# A maturation defective HIV-1 activates cGAS

**DOI:** 10.1101/2023.04.14.536845

**Authors:** Rebecca P. Sumner, Henry Blest, Meiyin Lin, Carlos Maluquer de Motes, Greg J. Towers

## Abstract

**Background:** Detection of viruses by host pattern recognition receptors induces the expression of type I interferon (IFN) and IFN-stimulated genes (ISGs), which suppress viral replication. Retroviruses such as HIV-1 are subject to sensing by both RNA and DNA sensors, and whether there are any particular features of the viral genome or reverse transcripts that facilitate or enhance this sensing is currently unknown.

**Results:** Whilst investigating the determinants of innate detection of HIV-1 we noticed that infection of THP-1 cells or primary macrophages with a virus expressing Gag fused to a reporter gene (luciferase or GFP) induced a robust IFN and ISG response that was not observed with an equivalent virus with similar genome length and composition, but expressing wild-type Gag. Innate immune activation by Gag-fusion HIV-1 was dependent on reverse transcription and DNA sensor cGAS, suggesting activation of an IFN response by viral DNA. Further investigation of the Gag-fusion viral particles revealed maturation defects, as evidenced by incomplete Gag cleavage and a diminished capacity to saturate restriction factor TRIM5α, likely due to aberrant particle formation. We propose that expression of the Gag fusion protein disturbs the correct cleavage and maturation of wild-type Gag, yielding viral particles that are unable to effectively shield viral DNA from detection by innate sensors including cGAS.

**Conclusions:** These data highlight the crucial role of capsid in innate evasion and support growing literature that disruption of Gag cleavage and capsid formation induces a viral DNA- and cGAS-dependent innate immune response. Together these data demonstrate a protective role for capsid and suggest that antiviral activity of capsid-targeting antivirals may benefit from enhanced innate and adaptive immunity *in vivo*.

## Background

Viral infection can be sensed by host pattern recognition receptors (PRRs) that detect viral nucleic acids and/or proteins. PRR engagement activates transcription factors belonging to the nuclear factor kappa-light-chain-enhancer of activated B cells (NF-κB) and interferon (IFN) regulatory factor (IRF) families, to induce expression of type I IFNs and inflammatory cytokines and chemokines[1]. IFNs activate signalling cascades dependent on Janus kinase (JAK) and signal transducer and activator of transcription (STAT) and the expression of IFN-stimulated genes (ISGs), including viral restriction factors[2]. A series of studies have demonstrated sensing of HIV-1 by RNA and DNA sensors. For example, the RNA genome has been reported to be sensed by DDX3[3] and MDA5[4] and viral DNA reverse transcripts by cyclic GMP-AMP synthase (cGAS)[5–7], IFI16[8, 9], PQBP1[10, 11] and NONO[12]. Further, DDX41 may sense RNA/DNA hybrids formed during reverse transcription[13]. Importantly, the central HIV DNA sensor appears to be cGAS, as it is required for HIV detection by other DNA sensors. cGAS is DNA sequence independent and when activated catalyses synthesis of cyclic GMP-AMP (2’,3’- cGAMP)[14–16] which induces STING phosphorylation and translocation to perinuclear regions [17]. STING recruitment of TBK1 and IRF3 results in IRF3 phosphorylation by TBK1 and IRF3 nuclear translocation[18, 19]. Activated STING also activates IKK and the NF-κB family of transcription factors[20], which with IRF3, activate expression of type I IFN and subsequently ISGs. ISGs include an array of anti-HIV restriction factors including APOBEC3G, SAMHD1, tetherin, TRIM5α, MxB and the IFITMs[21]. Despite all these examples of HIV-1 sensing, other studies demonstrate HIV replication in permissive primary cells without IFN induction. We hypothesise that sensing is context and particularly viral dose dependent. Thus whilst high dose infection can be sensed, particularly in cells that do not support HIV replication, e.g dendritic cells[6, 22], in permissive macrophages and T-cells, HIV-1 replication is a poor stimulator of IFN[23, 24] and the virus can replicate without triggering innate immune sensing through hiding nucleic acid PAMPs inside intact capsids[7, 25, 26], which uncoat and release genome inside the nucleus immediately prior to integration[27–30].

Growing evidence supports a crucial role for cellular cofactors in HIV-1 avoiding host immunity. Recruitment of cleavage and polyadenylation specificity factor 6 (CPSF6) and cyclophilin A (cypA) promote evasion of sensing, with cypA being particularly important for escaping HIV-1 capsid sensing by TRIM5α[7, 31]. Conversely, other cellular proteins that target the HIV-1 capsid, including NONO[12] and PQBP1[11], have been described to promote sensing by cGAS. In order to better understand the role of the HIV-1 capsid in sensing, and establish whether it promotes evasion, or is responsible for HIV-1 detection in infected cells, we tested the effect of making HIV-1 by co-expressing a truncated capsid with wild type Gag-pol. We found that truncated Gag fused to luciferase or GFP had a dominant negative effect on wild type Gag cleavage and caused a potent IFN response in THP-1 cells and macrophages that was not observed with wild-type (WT) HIV-1. Truncated Gag bearing viruses showed defective cleavage of wild type Gag, and failed to saturate TRIM5α or shield viral DNA from cGAS detection. These findings further evidence a role for the HIV-1 capsid in protecting HIV-1 genome from being sensed and support a model in which the principle function of capsid is to protect viral genomes from sensors to promote replication in sensing-competent target cells.

## Results

### HIV-1 Gag-fusion viruses trigger a robust type I IFN-dependent innate immune response in THP-1 cells

Whilst seeking to design an HIV-1 reporter by fusing luciferase (LUC) in frame with capsid (CA), we found that viruses made by co-transfecting the Gag-LUC reporter (Suppl Fig 1, Fig 1A) with wild type Gag-pol, and a VSV-G envelope, triggered sensing in THP-1 cells. The Gag-LUC reporter was based on HIV-1 LAI strain[32] and also encodes GFP in the place of Nef. It activated a dose-dependent innate immune response whilst, the WT VSV-G pseudotyped ΔEnv LAI-GFP did not, as previously observed [26] (LAI, Suppl Fig 1, Fig 1A). Innate induction was assessed by measuring luciferase activity in the supernatants of infected monocytic THP-1 cells that had been modified to express Gaussia luciferase under the control of the *IFIT-1* (also known as *ISG56*) promoter, which is both IRF-3- and IFN-sensitive[33]. Virus dose in these experiments was normalised according to RT activity, as measured by SG-PERT (see Methods). The number of infectious units per unit of RT (Suppl Fig 2A), or per genome copy (Suppl Fig 2B), was equivalent between WT and Gag-fusion viruses. Innate induction was not unique to the Gag-luciferase fusion as a second HIV-1 LAI virus carrying a similar Gag-GFP fusion also resulted in dose-dependent ISG induction (Gag-GFP, Suppl Fig 1, Fig 1A), ruling out an immunostimulatory feature in the luciferase sequence. Fusion of Gag to either GFP or luciferase makes it non-functional, therefore co-transfection with a WT Gag-pol packaging construct (e.g. p8.91, Suppl. Fig 1) is required to produce infectious particles. To rule out differences in 8.91 and LAI Gag sequences/proteins that could potentially explain the observed differences in innate immune activation, we also co-transfected WT LAI with 8.91 Gag-pol by co-transfecting the ΔEnv LAI genome and p8.91 packaging construct (8.91 LAI, Suppl Fig 1). This virus behaved the same as WT ΔEnv LAI alone and failed to induce ISG reporter activity at the doses tested, thus ruling out differences in Gag as an explanation for ISG induction in the Gag fusion viruses (Fig 1A).

**Figure 1:**
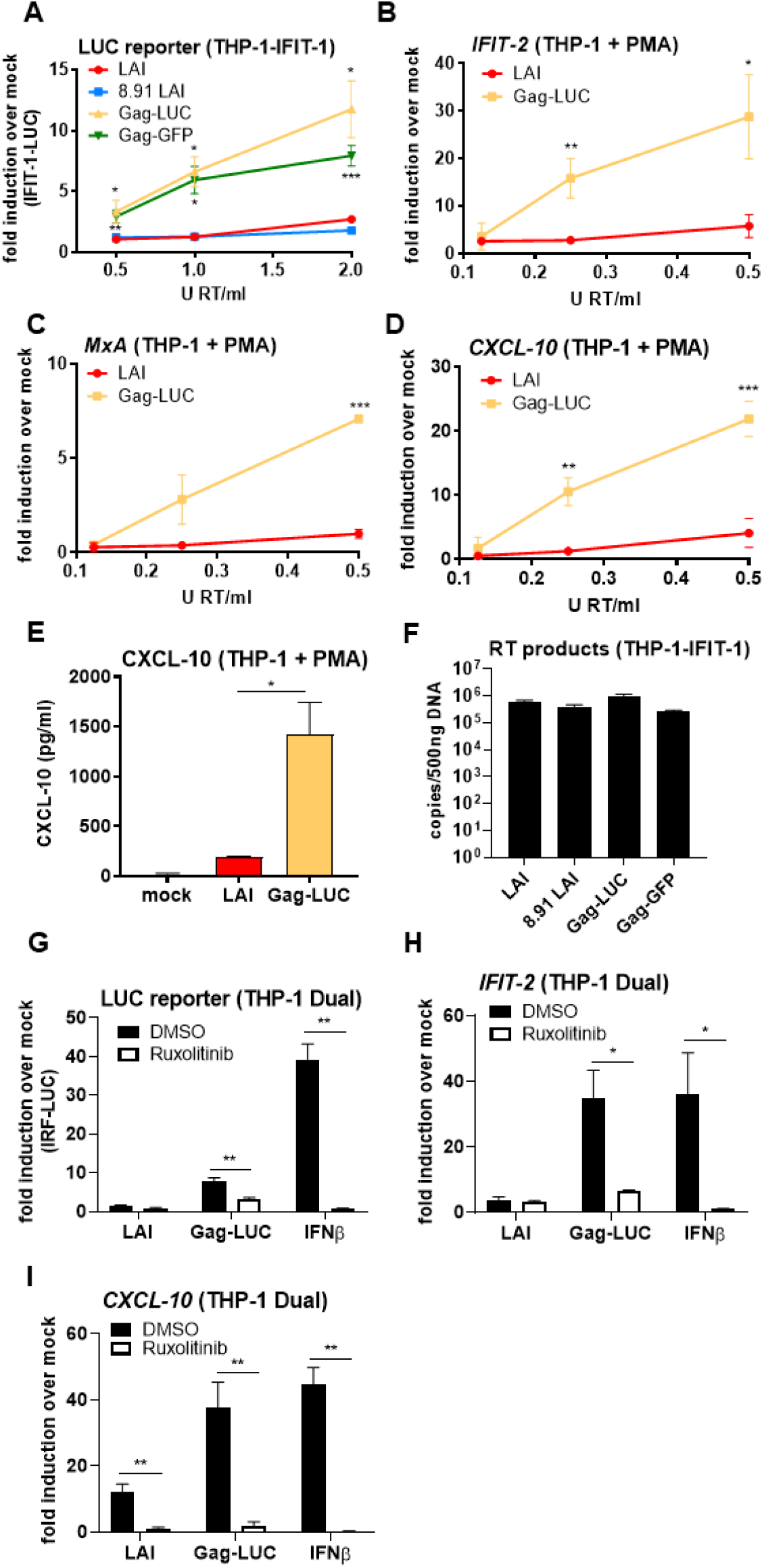
HIV-1 expressing a Gag-fusion protein triggers a type I IFN response in THP-1 cells A: IFIT-1 reporter activity from monocytic THP-1-IFIT-1 cells transduced for 24 h with WT LAI (LAI), LAI packaged with 8.91 Gag (8.91 LAI), LAI expressing gag fused to luciferase and packaged with 8.91 Gag (Gag-LUC) or LAI expressing Gag fused to GFP and packaged with 8.91 Gag (Gag-GFP) (See Suppl. Fig 1) at 0.5, 1 or 2 U RT/ml. B-D: ISG qRT-PCR from PMA-treated THP-1 shSAMHD1 cells transduced for 24 h with LAI or Gag-LUC viruses at 0.125, 0.25 and 0.5 U RT/ml. E: CXCL-10 protein in supernatants from B-D (0.5 U RT/ml, ELISA). F: RT products from THP-1-IFIT-1 cells transduced for 24 h with 1 U RT/ml of the indicated viruses. G: IRF reporter activity from monocytic THP-1 Dual cells transduced for 24 h with 1.5 U RT/ml LAI or Gag-LUC viruses, or stimulated with 1 ng/ml IFNβ as a control, in the presence of DMSO vehicle or 2 μM ruxolitinib. H, I: ISG qRT-PCR from monocytic THP-1 Dual cells transduced for 24 h with 1.5 U RT/ml LAI or Gag-LUC viruses, or stimulated with 1 ng/ml IFNβ as a control, in the presence of DMSO vehicle or 2 μM ruxolitinib. Data are mean ± SD, n = 3, representative of at least 3 repeats. Statistical analyses were performed using Student’s t-test, with Welch’s correction where appropriate, comparing each virus with WT LAI at the same dose (A-E) or pairs of samples −/+ ruxolitinib (G-I). *P < 0.05, **P < 0.01, ***P < 0.001.

To confirm the findings above from monocytic THP-1 cells, we also infected PMA differentiated THP-1 cells stably depleted for restriction factor SAMHD1. SAMHD1 was depleted to permit HIV transduction [26, 34]. The Gag-LUC virus, but not WT LAI, again induced high levels of endogenous ISGs *IFIT-2* (Fig 1B), *MxA* (Fig 1C) and *CXCL-10* (Fig 1D) measured by qPCR, as well as CXCL-10 protein (Fig 1E), measured by ELISA. Levels of viral reverse transcripts were equivalent in WT- and Gag fusion virus-infected cells, as assessed by qPCR (Fig 1F).

To assess whether Gag-fusion viruses induced type I IFN production we infected THP-1 Dual reporter cells (Invivogen) that also express luciferase under the control of an IRF- and ISG- sensitive promoter, in the presence of JAK1/2 inhibitor ruxolitinib[35]. Signal transduction downstream of the type I IFN receptor is dependent on JAK and thus ruxolitinib efficiently blocks IFNβ-induced ISG induction (Fig 1G-I). Expression of luciferase (Fig 1G), as well as endogenous *IFIT-2* (Fig 1H) and *CXCL-10* (Fig 1I) was significantly reduced following ruxolitinib treatment of Gag-LUC-infected cells indicating that infection with this Gag-LUC fusion virus induces type I IFN production, to induce endogenous ISG and IFN reporter expression.

### HIV-1 Gag-fusion viruses activate a restrictive type I IFN response in primary macrophages

To determine whether HIV-1 Gag-fusion viruses also induced a type I IFN response in primary human cells we infected primary monocyte-derived macrophages (MDM) with the Gag-LUC virus and the corresponding pseudotyped WT LAI strain and measured ISG induction by qPCR and ELISA. As in THP-1 cells, infection of MDM with Gag-LUC induced a robust type I IFN response leading to significantly higher expression of *CXCL-10* (Fig 2A), *IFIT-2* (Fig 2B) and *MxA* (Fig 2C), as well as CXCL-10 protein (Fig 2D) compared to VSV-G pseudotyped LAI infection, all of which was reduced by ruxolitinib treatment. Gag-LUC virus infection levels were lower in MDM than WT LAI at the same input dose, assessed by measuring GFP-positive cells by flow cytometry, and this was partially rescued by blocking IFN signalling with ruxolitinib indicating an IFN-dependent suppression of infection (Fig 2E). Taken together, Gag-fusion viruses, unlike their WT counterparts, induce a robust type I IFN response, which is restrictive even in a single round infection in primary macrophages.

**Fig 2:**
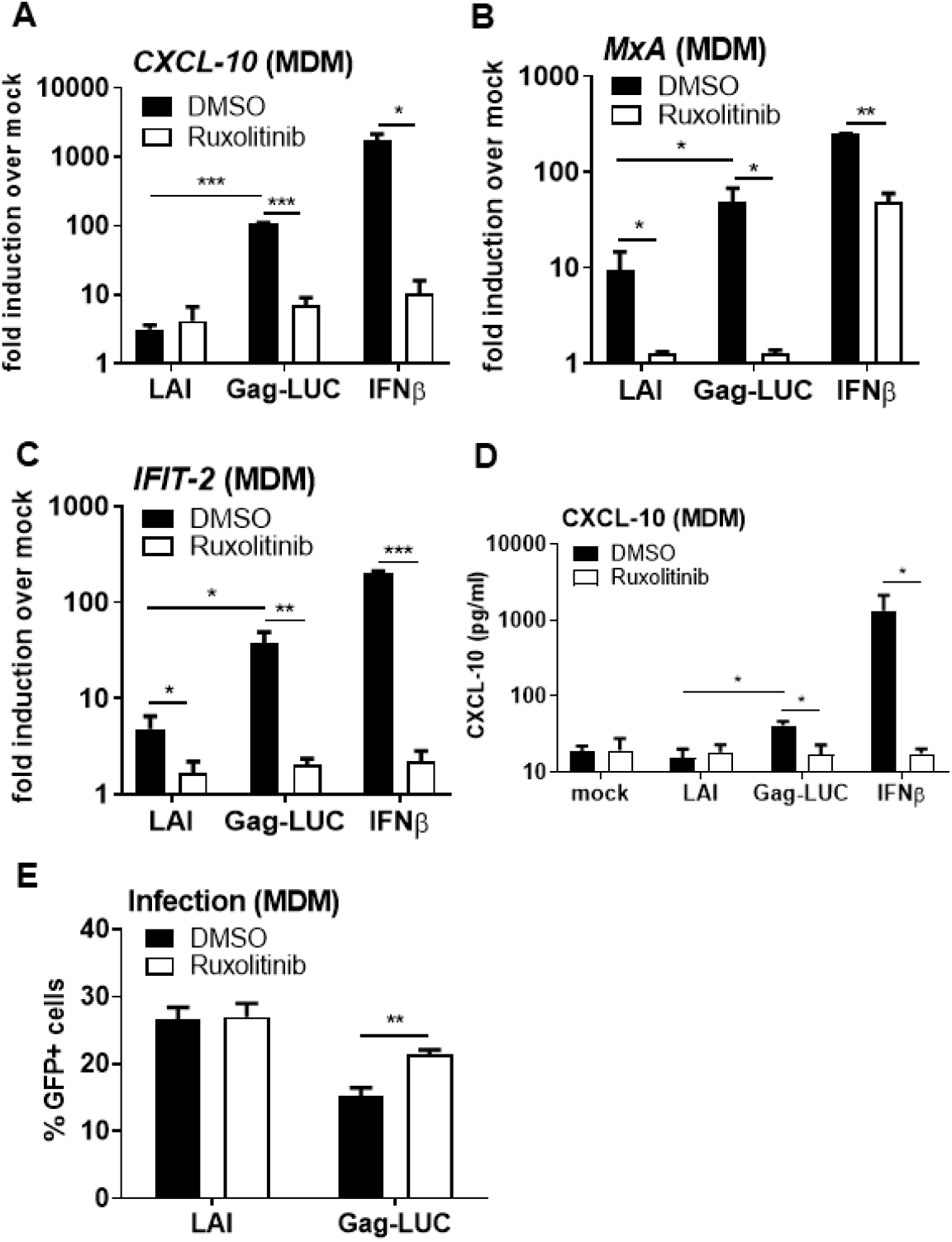
HIV-1 Gag-fusion viruses activate a restrictive type I IFN response in primary macrophages A-C: ISG qRT-PCR from primary MDM transduced for 24 h with 0.5 U RT/ml LAI or Gag-LUC viruses, or stimulated with 1 ng/ml IFNβ as a control, in the presence of DMSO vehicle or 2 μM ruxolitinib. D: CXCL-10 protein in supernatants from A-C (ELISA). E: Infection data from A-D measured by flow cytometry at 48 h. Data are mean ± SD, n = 3, representative of at least 3 repeats. Statistical analyses were performed using Student’s t-test, with Welch’s correction where appropriate, comparing pairs of samples −/+ ruxolitinib as indicated. *P < 0.05, **P < 0.01, ***P < 0.001.

### IFN induction by HIV-1 Gag-fusion viruses is dependent on viral DNA synthesis

To establish whether the source of immune stimulation during Gag-fusion virus infection was the viral genome, reverse transcripts, or a later stage of infection we generated Gag-LUC viruses that were defective for reverse transcription (Gag-LUC RT D185E) or integration (Gag- LUC INT D116N) by co-transfecting p8.91 Gag-pol carrying the RT D185E and INT D116N mutations. Luciferase IFN reporter in monocytic THP-1 IFIT-1 reporter cells (Fig 3A) and endogenous ISG induction in PMA-differentiated THP-1 shSAMHD1 cells (Fig 3B-D) was entirely RT-dependent (RT mutant did not trigger sensing) and did not require integration (Iitegrase mutant triggered normally). Concordantly, reporter activity (Fig 3E) and ISG expression (Fig 3F, G) was also significantly reduced in monocytic THP-1 Dual reporter cells following treatment with RT inhibitor nevirapine, but not with integrase inhibitor raltegravir.

**Fig 3:**
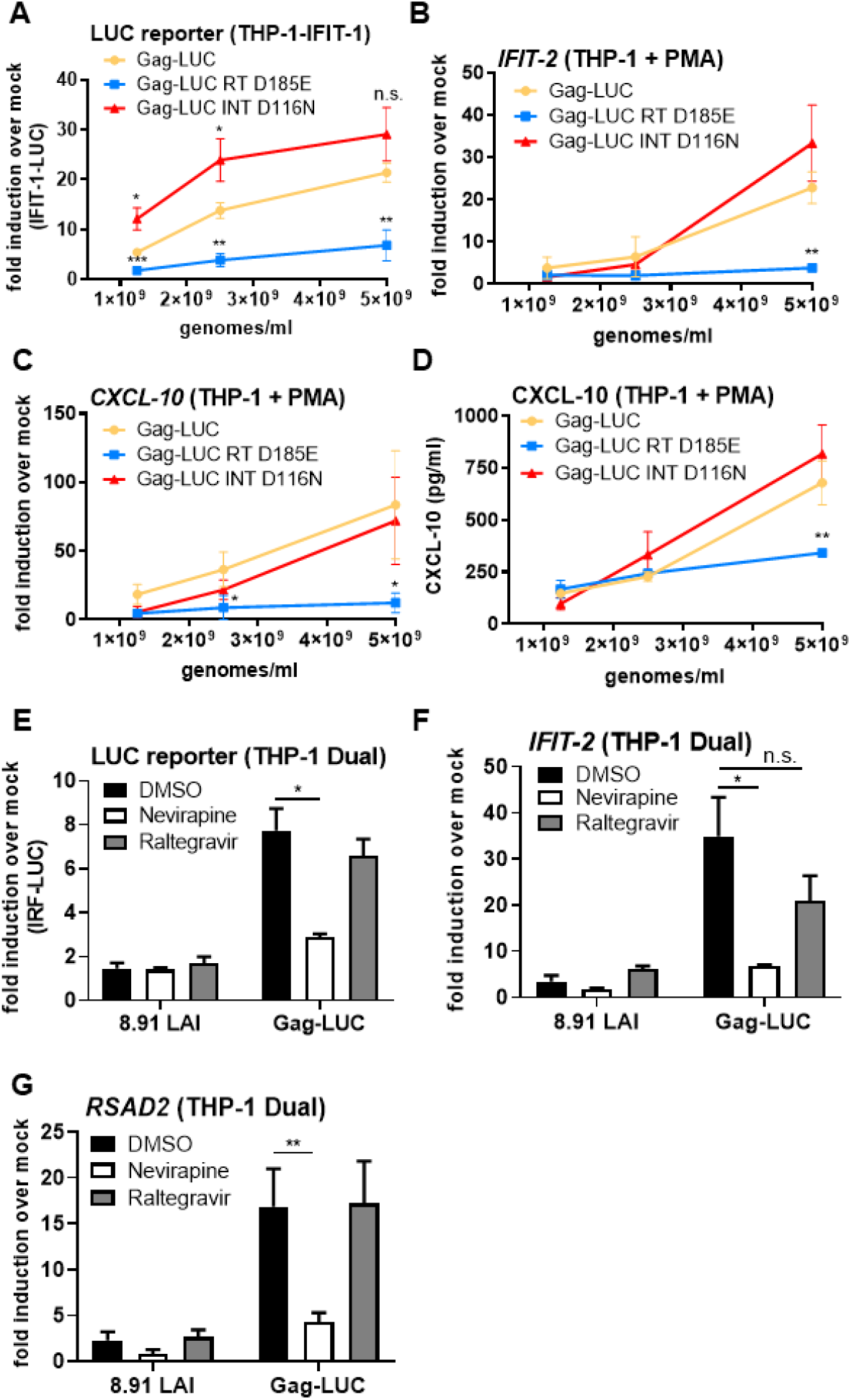
ISG induction by HIV-1 Gag-fusion virus is RT-dependent A: IFIT-1 reporter activity from monocytic THP-1-IFIT-1 cells transduced for 24 h with Gag-LUC, RT-defective Gag-LUC (Gag-LUC RT D185E) or integrase-defective Gag-LUC (Gag-LUC INT D116N) at 1.25×10^9^, 2.5×10^9^ and 5×10^9^ genomes/ml. B, C: ISG qRT-PCR from PMA-treated THP-1 shSAMHD1 cells transduced for 24 h with Gag- LUC, Gag-LUC RT D185E or Gag-LUC INT D116N at 1.25×10^9^, 2.5×10^9^ and 5×10^9^ genomes/ml. D: CXCL-10 protein in supernatants from B, C (ELISA). E: IRF reporter activity from THP-1 Dual cells transduced for 24 h with 8.91 LAI or Gag-Luc (1.5 U RT/ml) in the presence of DMSO vehicle, 5 μM neviripine or 10 μM raltegravir. F, G: ISG qRT-PCR from THP-1 Dual cells transduced for 24 h with 8.91 LAI or Gag-Luc (1.5 U RT/ml) in the presence of DMSO vehicle, 5 μM neviripine or 10 μM raltegravir. Data are mean ± SD, n = 3, representative of at least 3 repeats. Statistical analyses were performed using Student’s t-test, with Welch’s correction where appropriate, comparing mutant viruses with WT Gag-LUC at the same dose (A-D) or to the DMSO control as indicated (E-G). *P < 0.05, **P < 0.01, n.s. non-significant.

As expected, no GFP positive cells were observed following Gag-LUC RT D185E infection (Suppl Fig 3A, B) and levels of infectivity were also significantly reduced following nevirapine treatment (Suppl Fig 3C). Whilst GFP positivity was minimal in integrase defective Gag-LUC infection in differentiated THP-1 cells (Suppl Fig 3B), GFP positive cells were still detected with Gag-LUC INT D116N infection (Suppl Fig 3A) or following raltegravir treatment (Suppl Fig 3C) in monocytic THP-1 cells. This is in agreement with our previous findings[26] and likely due to GFP expression from unintegrated 2’-LTR circles that have been observed in other cell types[36, 37]. Together, these data rule out the viral RNA genome as the immunostimulatory feature of the Gag-fusion viruses and instead point to innate immune detection of viral DNA.

### ISG induction by HIV-1 Gag-fusion virus is dependent on cGAS and STING

To further investigate the source for immune stimulation in the Gag-fusion viruses we sought to determine which host innate sensors were required for innate immune detection. As expected, THP-1 IFIT-1 reporter cells lacking STING failed to respond to herring testis DNA (HT-DNA) stimulation, but did respond to transfected RNA mimic poly I:C and TLR4 agonist lipopolysaccharide (LPS). MAVS -/- cells responded to HT-DNA and LPS, but not transfected poly I:C (Suppl. Fig 4A). Luciferase reporter activity (Fig 4A) and endogenous ISG expression (Fig 4B, C) of Gag-LUC infection was entirely dependent on STING. Levels of infection were equivalent between WT and STING- or MAVS-null cells (Suppl Fig 4B). Furthermore, THP-1 Dual cells lacking cGAS failed to respond to HT-DNA (Suppl Fig 4C) and Gag-LUC infection (Fig 4D-F), consistent with a cGAS/STING-dependent DNA sensing response. Again, levels of infection were equivalent in WT and cGAS-/- cells (Suppl Fig 4D). Finally luciferase reporter activity in Gag-LUC infected THP-1 Dual cells was significantly reduced in the presence of STING inhibitor H151[38] and cGAS inhibitor RU.521[39] (Fig 4G, Suppl Fig 4E), confirming cGAS/STING-dependent sensing of viral reverse transcripts during Gag-fusion virus infection.

**Fig 4:**
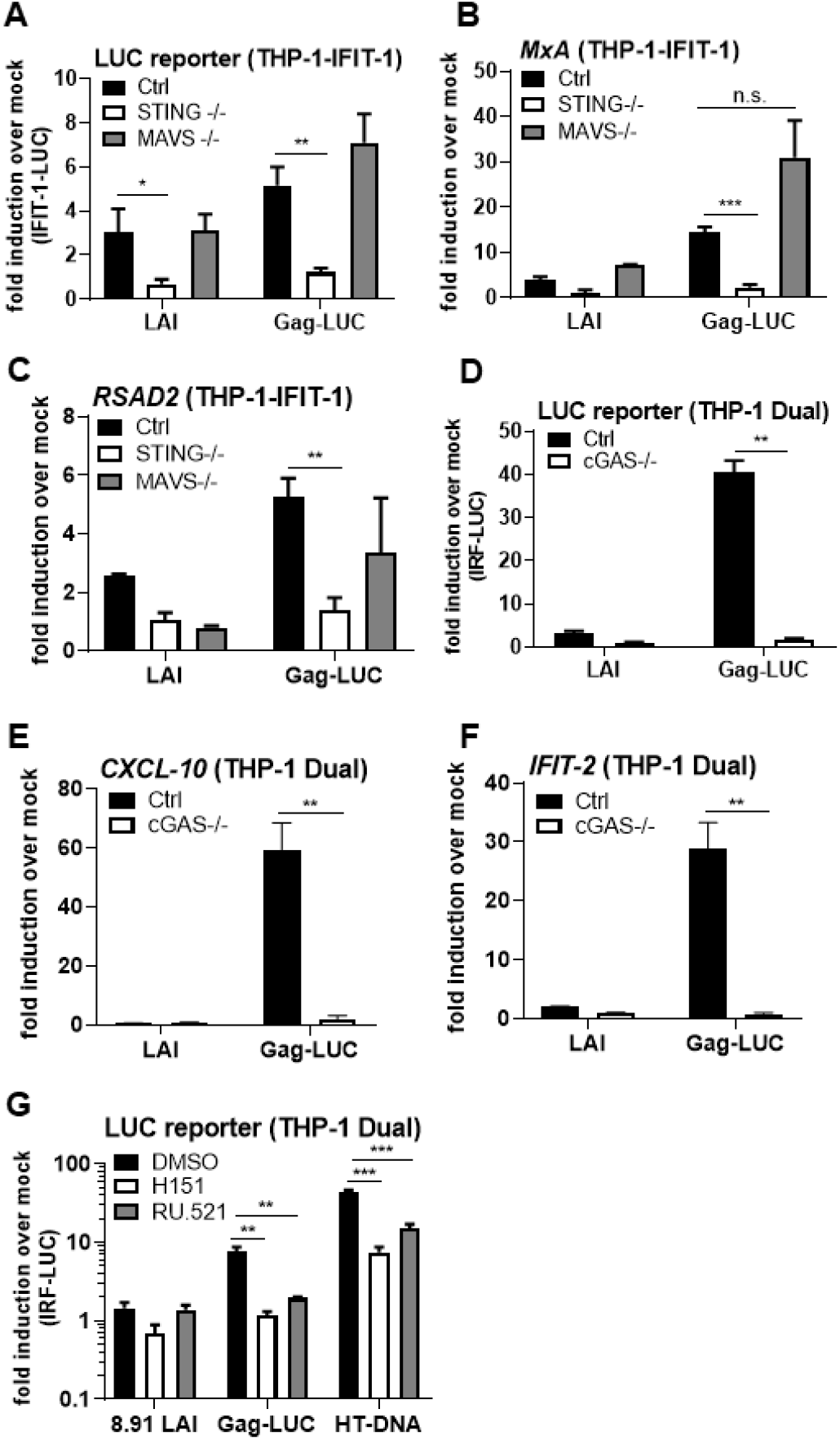
ISG induction by HIV-1 Gag-fusion virus is dependent on cGAS and STING A: IFIT-1 reporter activity from monocytic THP-1-IFIT-1 cells lacking STING or MAVS, or a gRNA control (Ctrl) cell line transduced for 24 h with WT LAI or Gag-LUC (1.5 U RT/ml). B, C: ISG qPCR from monocytic THP-1-IFIT-1 cells lacking STING or MAVS, or a gRNA control (Ctrl) cell line transduced for 24 h with WT LAI or Gag-LUC (1.5 U RT/ml). D: IRF reporter activity from monocytic THP-1 Dual cells lacking cGAS, or a gRNA control (Ctrl) cell line transduced for 24 h with WT LAI or Gag-LUC (1.5 U RT/ml). E, F: ISG qPCR from monocytic THP-1 Dual cells lacking cGAS, or a gRNA control (Ctrl) cell line transduced for 24 h with WT LAI or Gag-LUC (1.5 U RT/ml). G: IRF reporter activity from monocytic THP-1 Dual cells lacking cGAS, or a gRNA control (Ctrl) cell line transduced for 24 h with WT LAI or Gag-LUC (1.5 U RT/ml), or stimulated by transfection with 0.05 μg/ml HT-DNA in the presence of DMSO vehicle, 0.5 μg/ml STING inhibitor H151 or 10 μg/ml cGAS inhibitor RU.521. Data are mean ± SD, n = 3, representative of at least 3 repeats. Statistical analyses were performed using Student’s t-test, with Welch’s correction where appropriate, comparing to Ctrl cells (A-F), or to DMSO vehicle treated cells (G) as indicated. *P < 0.05, **P < 0.01, ***P < 0.001, n.s. non-significant.

### Gag-fusion viruses display defects in maturation and are less able to saturate TRIM5α

Given that the genome sequences of the LAI and Gag-LUC/Gag-GFP viruses only differ by the inclusion of the chimeric Gag-LUC/GFP reporter gene, and encode for all the same accessory proteins, we hypothesised that rather than specific features of the genome enhancing sensing, the Gag-fusion viruses may instead fail to efficiently shield RT products from cGAS through a dominant negative effect of the Gag fusion. Indeed, immunoblots of extracted viral particles, detecting HIV-1 capsid protein, showed that both Gag-fusion viruses had evidence of maturation defects, with increased levels of MA-NC and other partial Gag cleavage products below MA-CA (Fig 5A).

**Fig 5:**
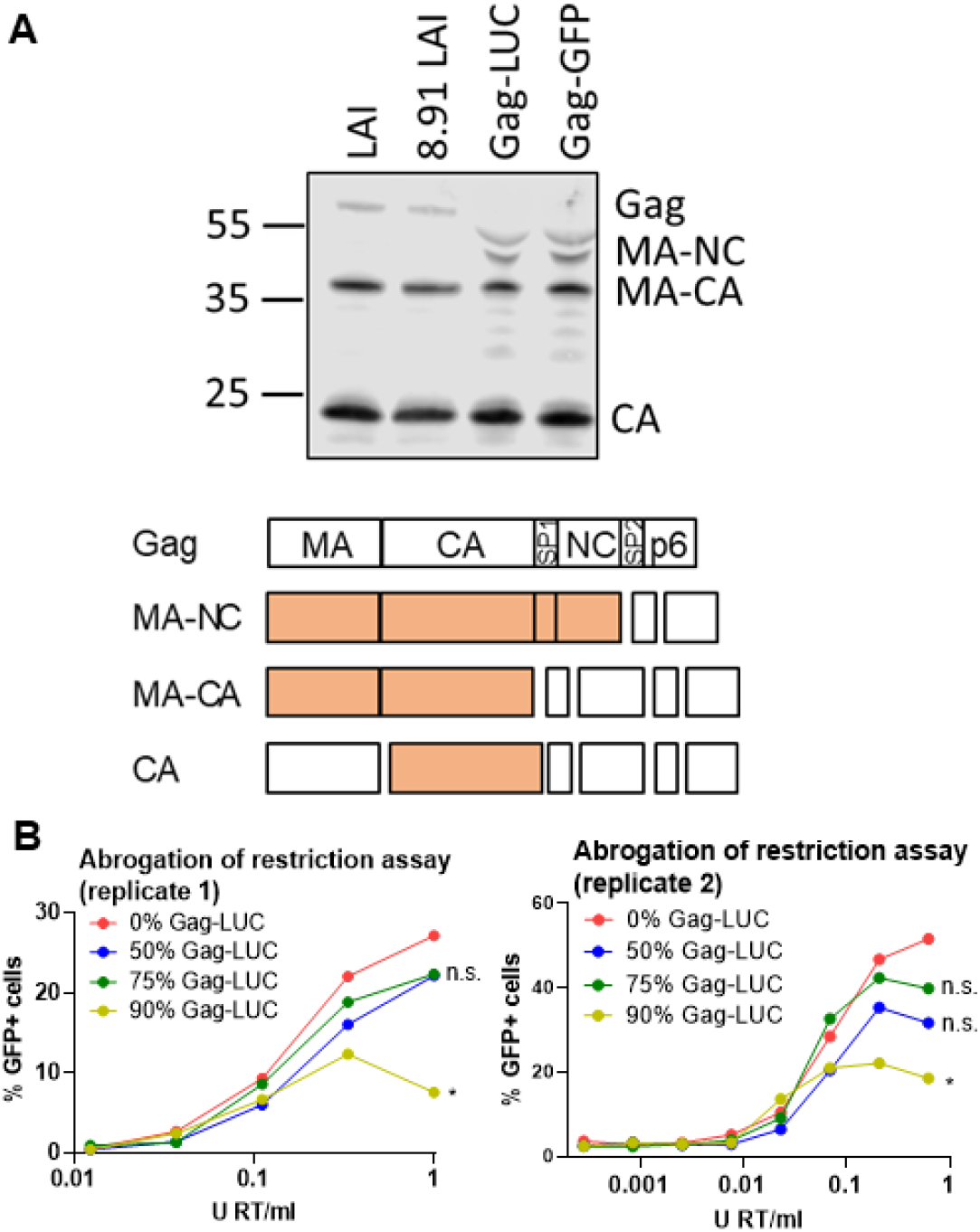
Gag-fusion viruses display defects in maturation and are less able to saturate TRIM5α A: Immunoblot of WT LAI, 8.91 LAI, Gag-LUC and Gag-GFP virus particles (2×10^11^ genomes) detecting p24 and a schematic of intermediate Gag cleavage products. MA: matrix, CA: capsid, SP1: spacer peptide 1, NC: nucleocapsid, SP2: spacer peptide 2. B: Abrogation-of-restriction assay in FRhK4 cells expressing restrictive rhesus TRIM5. FRhK4 cells were co-transduced with a fixed dose of WT LAI.GFP (5×10^7^ genomes/ml) and increasing doses of the WT/Gag-LUC chimeric viruses carrying a luciferase-expressing genome (0.0005 – 1 U RT/ml). Rescue of GFP infectivity was assessed by flow cytometry at 48 h. Data are singlet % GFP values and two repeats of the experiment are shown. Statistical analyses were performed using 2-way ANOVA with multiple comparisons. * *P*<0.05, n.s. non-significant.

To assess HIV-1 core integrity in the Gag-fusion viruses we measured their ability to saturate rhesus monkey TRIM5α in an abrogation-of-restriction assay. Rhesus monkey TRIM5α binds and forms hexameric cage-like structures around intact HIV capsid lattices[40, 41], leading to proteasome-dependent viral disassembly and subsequent innate immune activation[42–44]. Restriction by TRIM5α can be overcome by co-infection with high doses of a saturating virus, dependent on the stability of the incoming viral capsid[45, 46]. The Gag-LUC fusion protein was cloned into the p8.91 Gag-pol packaging plasmid and HEK 293T cells were transfected with varying proportions of WT or Gag-LUC p8.91, thus producing VSV-G psuedotyped viruses with increasing amounts of Gag-fusion protein. In all cases the same genome expressing luciferase (CSLW) was packaged. Rhesus FRhK cells were then co-infected with a fixed dose of HIV-1 LAI bearing GFP and increasing doses of the WT/Gag-LUC chimeric viruses. Flow cytometry was used to assess rescue of HIV-1 LAI infectivity from TRIM5α restriction measuring GFP positive cells. As expected, the virus with 100% WT Gag (0% Gag-LUC) efficiently saturated TRIM5α restriction and rescued HIV-1 LAI GFP expression (Fig 5B, Suppl. Fig 5). Increasing the proportion of luciferase-fused Gag in the saturating virus, reduced rescue of GFP expression, which reached statistical significance at the highest proportion of Gag-LUC (90% Gag-LUC, Fig 5B).

We conclude that expression of this Gag-LUC fusion protein during viral production interferes with the maturation process of co-transfected WT Gag, yielding particles with reduced stability and a diminished ability to saturate TRIM5α, which fail to shield their RT products from DNA sensor cGAS. This finding adds to growing literature that intact capsid plays a crucial role in HIV-1 evasion of cGAS and that antiviral activity of capsid-targeting antivirals may benefit from triggering innate immune detection and subsequent antiviral gene expression *in vivo*.

## Discussion

Numerous studies have described HIV-1 as a poor activator of innate immunity *in vitro*[6, 7, 23, 26] unless infection is high dose or target cells are not usually permissive to HIV replication e.g dendritic cells[6, 22]. This suggests that, like many other viruses, HIV-1 has evolved strategies to evade the host response. In addition to encoding accessory proteins that block innate signalling cascades and activation of transcription factors such as NF-κB and IRF3[47–52], growing evidence points to a critical role for capsid in innate immune evasion. Cellular cofactors CPSF6 and cyclophilin A are recruited by capsid and are critical for evasion of sensing, the latter being important for avoiding TRIM5α restriction[7, 31]. Encapsidated DNA synthesis is expected to protect viral RT products from DNA sensors such as cGAS and from degradation by cellular nucleases such as TREX-1[7, 45, 53]. Supporting this, recent studies have linked capsid stability to activation of cGAS sensing [54], including our own work demonstrating that disrupting capsid maturation using protease inhibitors, or by mutating cleavage sites in Gag, yields aberrant viral particles that fail to protect RT products from cGAS[26]. Furthermore, differences in the ability of HIV-1, and HIV-2 and other non-pandemic lentiviruses to evade innate immunity has been mapped to the viral capsid, with the ability to evade cGAS activation and TRIM5α correlating with pandemicity [6, 25]. In this study we report the unexpected finding that unlike WT HIV-1, HIV-1 viruses carrying a truncated Gag fusion protein trigger a robust type I IFN response in macrophages (Fig 1, 2), dependent on reverse transcription (Fig 3) and host DNA sensing machinery cGAS and STING (Fig 4). Importantly, virus made with Gag fusions showed evidence of maturation defects and had a reduced capacity to saturate restriction factor TRIM5α in an abrogation-of-restriction assay, indicative of defective capsids (Fig 5). This work adds to a growing body of evidence that the HIV-1 capsid plays a crucial role in shielding RT products from cGAS.

Exactly how the expression of Gag fused to a reporter gene such as luciferase or GFP inhibits Gag cleavage and functional capsid formation is not known. Given maturation occurs post-budding it is plausible that the Gag fusion proteins are incorporated into nascent virions and interfere with maturation. The defective viruses may have altered stability, may prematurely uncoat and subsequently activate a potent host innate response that is not observed for similar doses of WT virus. Furthermore, interactions with host proteins may also differ, and whether the Gag-LUC and Gag-GFP viruses used in this study still interact appropriately with cofactors including as CPSF6 and cypA, or incorporate the capsid stabilising cellular metabolite inositol hexakisphosphate (IP6) that is dependent on the immature lattice[55], remains to be determined.

Thus far cGAS has been described to sense DNA in a sequence-independent manner[15, 56], but whether there are particular features of viruses or their genomes that enhance recognition is unclear. Additional proteins may be involved in fine-tuning the cGAS response or breaking capsid open to expose viral DNA within. For example, PQBP1 has recently been described to directly bind and decorate the HIV-1 capsid, ‘licensing’ it for subsequent cGAS recruitment and sensing of viral DNA[11].

As we have previously observed[26], activation of an IFN response by maturation defective viruses during single round infection of THP-1 cells was not sufficient to block infection, with WT and Gag-LUC/GFP viruses being equally infectious in THP-1 and U87 cells (Suppl Fig 2). Infectivity of the Gag-LUC virus was however reduced compared to WT in primary macrophages, and this was partially rescued by blocking IFN signalling (Suppl Fig 2E). Primary cells may express higher levels of IFN, be more sensitive to IFN, or may express a wider range of restrictive ISGs than cell lines such as THP-1 that could explain these differences. Unprotected RT products during Gag-LUC infection may also be subject to degradation by TREX1, which could also account for some of the remaining restriction in MDM.

In summary we have discovered an unanticipated effect on the maturation of WT Gag by coexpression of a Gag fusion protein, yielding aberrant viral particles that fail to shield their DNA from cGAS and induce a restrictive type I IFN response in macrophages. This finding supports the crucial role of capsid in innate immune evasion and highlights this viral protein as an important target for novel therapeutics. Indeed, it will be interesting to test whether recently described capsid-targeting inhibitors, such as those from Gilead [57], also induce sensing of HIV-1 RT products as we recently demonstrated for PF-74[26], which accelerates capsid opening[58]. Likewise, maturation inhibitors such as bevirimat[59] may also lead to enhanced sensing in a similar manner to that observed with protease inhibitors[26]. It remains to be seen whether capsid or protease inhibitors leverage innate immune responses to improve their efficacy *in vivo*.

## Materials and Methods

### Cells and reagents

HEK293T, FRhK and U87 cells were maintained in DMEM (Gibco) supplemented with 10 % foetal bovine serum (FBS, Labtech) and 100 U/ml penicillin plus 100 μg/ml streptomycin (Pen/Strep; Gibco). THP-1 cells were maintained in RPMI (Gibco) supplemented with 10 % FBS and Pen/Strep. THP-1-IFIT-1 cells that had been modified to express Gaussia luciferase under the control of the *IFIT-1* promoter[33] and versions lacking MAVS or STING[34] were described previously. THP-1 cells stably depleted for SAMHD1 were also previously described[26]. THP-1 Dual Control and cGAS-/- cells were obtained from Invivogen. Nevirapine and raltegravir were obtained from AIDS reagents. STING inhibitor H151 and cGAS inhibitor RU.521 were obtained from Invivogen. JAK inhibitor ruxolitinib was obtained from CELL guidance systems. Lipopolysaccharide and IFNβ were obtained from Peprotech. Herring-testis DNA was obtained from Sigma. cGAMP and poly I:C were obtained from Invivogen. For stimulation of cells by transfection, transfection mixes were prepared using lipofectamine 2000 according to the manufacturer’s instructions (Invitrogen).

### Isolation of primary monocyte-derived macrophages

Primary monocyte-derived macrophages (MDM) were prepared from fresh blood from healthy volunteers as described previously[26]. The study was approved by the joint University College London/University College London Hospitals NHS Trust Human Research Ethics Committee and written informed consent was obtained from all participants. Replicate experiments were performed with cells derived from different donors.

### Generation of Gag fusion, RT D185E and INT D116N viruses

pLAIΔEnvGFP.Gag-LUC/GFP and p8.91 Gag-LUC were generated by cloning the firefly luciferase gene/GFP into the unique SpeI site of CA. pLAIΔEnvGFP.Gag-LUC RT D185E and INT D116N were generated by site-directed mutagenesis using Pfu Turbo DNA polymerase (Agilent) and the following primers:

LAI_ RT D185E fwd: 5’ ATAGTTATCTATCAATACATGGAAGATTTGTATG 3’

LAI_ RT D185E rev: 5’ AAGTCAGATCCTACATACAAATCTTCCATGTATTG 3’

LAI_ INT D116N fwd: 5’ GGCCAGTAAAAACAATACATACAAACAATGGCAGC 3’

LAI_ INT D116N rev: 5’ ACTGGTGAAATTGCTGCCATTGTTTGTATGTATTG 3’

In all cases mutated sequences were confirmed by sequencing, excised by restriction digestion and cloned back into the original plasmid.

### Viral production in HEK293T cells

Lentiviral particles were produced by transfection of HEK293T cells in T150 flasks using Fugene 6 transfection reagent (Promega) according to the manufacturer’s instructions. For LAI WT each flask was transfected with 2.5 μg of VSV-G glycoprotein expressing plasmid pMDG (Genscript) and 6.25 μg pLAIΔEnvGFP (Suppl. Fig. 1). For viruses requiring a packaging plasmid each flask was transfected with 2.5 μg of pMDG (Genscript), 2.5 μg of p8.91 (encoding Gag-Pol, Tat and Rev)[60], and 3.75 μg of genome plasmid (pLAIΔEnvGFP, pLAIΔEnvGFP.Gag-LUC, pLAIΔEnvGFP.Gag-GFP, Suppl. Fig. 1). WT/Gag-LUC chimeric viruses were generated by transfecting cells with 2.5 μg of pMDG, 3.75 μg of a firefly luciferase-expressing genome plasmid (CSLW) and varying proportions of p8.91 and p8.91Gag-LUC packaging plasmids, up to 2.5 μg per flask. Virus supernatants were harvested at 48 and 72 h post-transfection, pooled, DNase treated (2 h at 37 °C, DNaseI, Sigma) and subjected to ultracentrifugation over a 20 % sucrose cushion. Viral particles were resuspended in RPMI supplemented with 10 % FBS. Viral titres were calculated by infecting PMA-treated THP-1 cells (2×10^5^ cells/ml) or U87 cells (10^5^ cells/ml) with dilutions of virus in the presence of polybrene (8 μg/ml, Sigma) for 48 h and enumerating GFP-positive cells by flow cytometry using the FACS Calibur (BD). Analysis was performed using FlowJo software.

### SG-PERT

Reverse transcriptase activity of virus preparations was quantified by qPCR using a SYBR Green-based product-enhanced RT (SG-PERT) assay as described [61].

### Genome copy/RT products measurements

Viral genome copies and RT products were measured by qPCR as previously described using primers specific for GFP[26]:

*GFP* fwd: 5’- CAACAGCCACAACGTCTATATCAT -3’

*GFP* rev: 5’- ATGTTGTGGCGGATCTTGAAG -3’

*GFP* probe: 5’- FAM-CCGACAAGCAGAAGAACGGCATCAA-TAMRA -3’

### Infection assays

THP-1 cells were infected at a density of 2×10^5^ cells/ml in 24 well plates for luciferase reporter assays or 12 well plates for qPCR and ELISA. For differentiation, THP-1 cells were treated with 50 ng/ml phorbol 12-myristate 13-acetate (PMA, Peprotech) for 48 h. Infections in THP-1 cells were performed in the presence of polybrene (8 μg/ml, Sigma). Input dose of virus was normalised either by RT activity (measured by SG-PERT) or genome copies (measured by qPCR) as indicated. Infection levels were assessed at 48 h post-infection through enumeration of GFP positive cells by flow cytometry.

### Luciferase reporter assays

Gaussia/Lucia luciferase activity in supernatants was measured by transferring 10 μl to a white

96 well assay plate, injecting 50 μl per well of coelenterazine substrate (Nanolight Technologies, 2 μg/ml) and analysing luminescence on a FLUOstar OPTIMA luminometer (Promega). Fold inductions were calculated by normalising to a mock-treated control.

### ISG qPCR

ISG induction in infected THP-1 cells and primary MDM was assessed by qPCR as previously described[26]. Expression of each gene was normalised to an internal control (*GAPDH*) and these values were then normalised to mock-treated control cells to yield a fold induction. The following primers were used:

*GAPDH* Fwd: 5’-GGGAAACTGTGGCGTGAT-3’,

*GAPDH* Rev: 5’-GGAGGAGTGGGTGTCGCTGTT-3’

*CXCL-10* Fwd: 5’-TGGCATTCAAGGAGTACCTC-3’

*CXCL-10* Rev: 5’-TTGTAGCAATGATCTCAACACG-3’

*IFIT-2* Fwd: 5’-CAGCTGAGAATTGCACTGCAA-3’

*IFIT-2* Rev: 5’-CGTAGGCTGCTCTCCAAGGA-3’

*MxA* Fwd: 5’-ATCCTGGGATTTTGGGGCTT-3’

*MxA* Rev: 5’-CCGCTTGTCGCTGGTGTCG-3’

*RSAD2* Fwd: 5’-CTGTCCGCTGGAAAGTG-3’

*RSAD2* Rev: 5’-GCTTCTTCTACACCAACATCC-3’

### ELISA

Cell supernatants were harvested for ELISA at 24 h post-infection/stimulation and stored at −80 °C. CXCL-10 protein was measured using Duoset ELISA reagents (R&D Biosystems) according to the manufacturer’s instructions.

### Immunoblotting

For immunoblotting of viral particles, 2×10^11^ genome copies of virus were boiled for 10 min in 6X Laemmli buffer (50 mM Tris-HCl (pH 6.8), 2 % (w/v) SDS, 10% (v/v) glycerol, 0.1% (w/v) bromophenol blue, 100 mM β-mercaptoethanol) before separating on 4-12 % Bis-Tris polyacrylamide gradient gel (Invitrogen). After PAGE, proteins were transferred to a Hybond ECL membrane (Amersham biosciences) using a semi-dry transfer system (Biorad). Mouse- anti-HIV-1capsid p24 was from AIDS reagents (183-H12-5C) and was detected with goat-anti- mouse IRdye 800CW infrared dye secondary antibody and membranes imaged using an Odyssey Infrared Imager (LI-COR Biosciences).

### Abrogation-of-restriction assay

FRhK cells were plated in 48 well plates at 5×10^4^ cells/ml. The following day cells were co- transduced in the presence of polybrene (8 μg/ml, Sigma) with a fixed dose of HIV-1 LAI expressing GFP (5×10^7^ genome copies/ml) and increasing doses of the WT/Gag-LUC chimeric viruses carrying a luciferase-expressing genome, CSLW (0.0005 – 1 U RT/ml). Rescue of GFP infectivity was assessed 48 h later by flow cytometry using the FACS Calibur (BD) and analysing with FlowJo software.

### Statistical analyses

Statistical analyses were performed using an unpaired Student’s t-test (with Welch’s correction where variances were unequal) or a 2-way ANOVA with multiple comparisons, as indicated. * *P*<0.05, ** *P*<0.01, *** *P*<0.001.

## List of abbreviations

cGAMP: cyclic GMP-AMP
cGAS: cyclic GMP-AMP synthase
CA: capsid
CPSF6: cleavage and polyadenylation specificity factor 6
cypA: cyclophilin A
env: envelope
GFP: green fluorescent protein
HIV: human immunodeficiency virus
HT-DNA: herring testis DNA
IFN: interferon
IP6: inositol hexakisphosphate 6
IRF: interferon regulatory factor
ISG: interferon stimulated gene
IU: infectious unit
JAK: Janus kinase
LPS: lipopolysaccharide
LTR: long terminal repeat
Luc: luciferase
MA: matrix
MDM: monocyte-derived macrophage
NC: nucleocapsid
NF-𝜅B: nuclear factor kappa B
PAMP: pathogen-associated molecular pattern
PMA: phorbol 12-myristate 13-acetate
PRR: pattern recognition receptor
RT: reverse transcriptase
SG-PERT: SYBR Green-based product-enhanced
RT SP: spacer peptide
STAT: signal transducer and activator of transcription
WT: wild-type
VSV-G: vesicular stomatitis virus G protein

## Declarations

### Ethics approval

Not applicable.

### Consent for publication

Not applicable.

### Availability of data and materials

All data generated or analysed during this study are included in this published article [and its supplementary information files].

### Authors’ Contributions

RPS and GJT conceptualised the study. RPS, HB and ML performed the experiments and analysed the data. RPS, CMM and GJT wrote the manuscript. GJT and CMM obtained funding. All authors read and approved the final manuscript.

## Acknowledgements

We thank Veit Hornung for providing THP-1-IFIT-1 cells.

---

**Suppl Fig 1:**
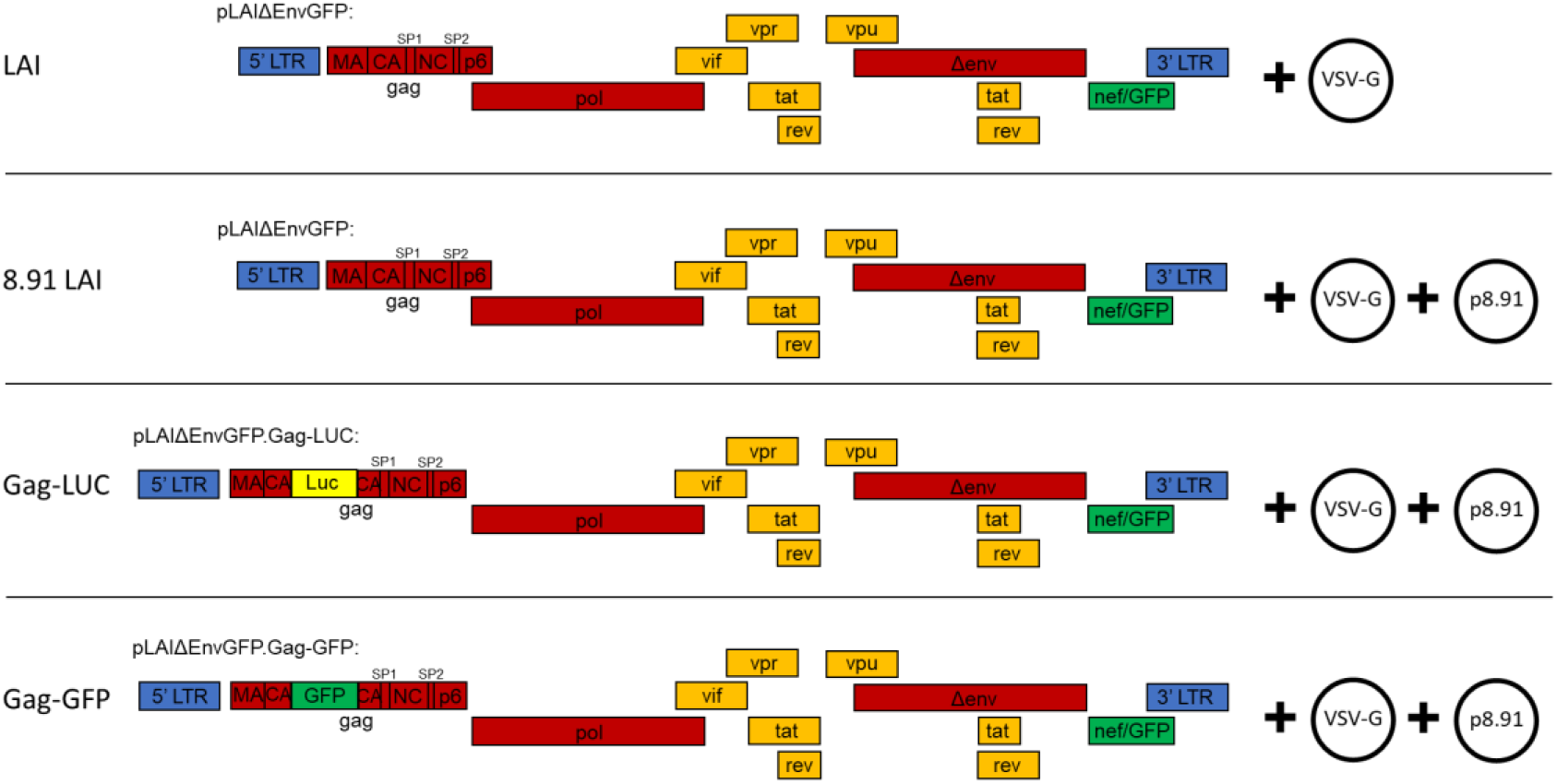
Schematic of wild-type and Gag-fusion viruses Schematic representation of the plasmids transfected into HEK293T cells to produce WT and Gag-fusion viruses. The genome plasmid of each virus is based on the HIV-1 LAI strain[32] with a deletion in envelope (Δenv) and expressing GFP in the place of Nef (pLAIΔEnvGFP). Each virus was pseudotyped with VSV-G, and for 8.91 LAI, Gag-LUC and Gag-GFP viruses, were co-transfected with p8.91 packaging construct encoding Gag-Pol, Tat and Rev[60]. LTR: long terminal repeat, MA: matrix, CA: capsid, SP: spacer peptide, NC: nucleocapsid, env: envelope, Luc: firefly luciferase, GFP: green fluorescent protein.

**Suppl Fig 2:**
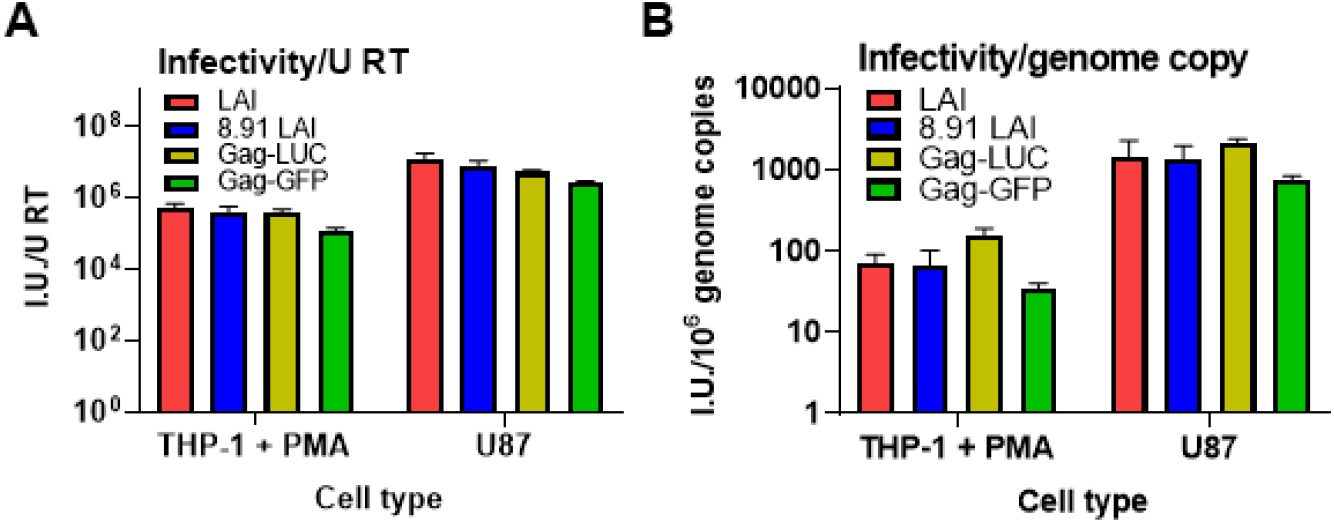
Particle infectivity of wild-type and Gag-fusion viruses A: Virus infectivity (infectious units, I.U.) on THP-1 cells differentiated with PMA or U87 cells (measured by flow cytometry at 48 h post-transduction) normalised to units of RT (measured by SG-PERT). B: Virus infectivity (infectious units, I.U.) on THP-1 cells differentiated with PMA or U87 cells (measured by flow cytometry at 48 h post-transduction) normalised to genome copy number (measured by qPCR). Data are mean ± SD, n = 3, representative of at least 2 repeats.

**Suppl. Fig 3:**
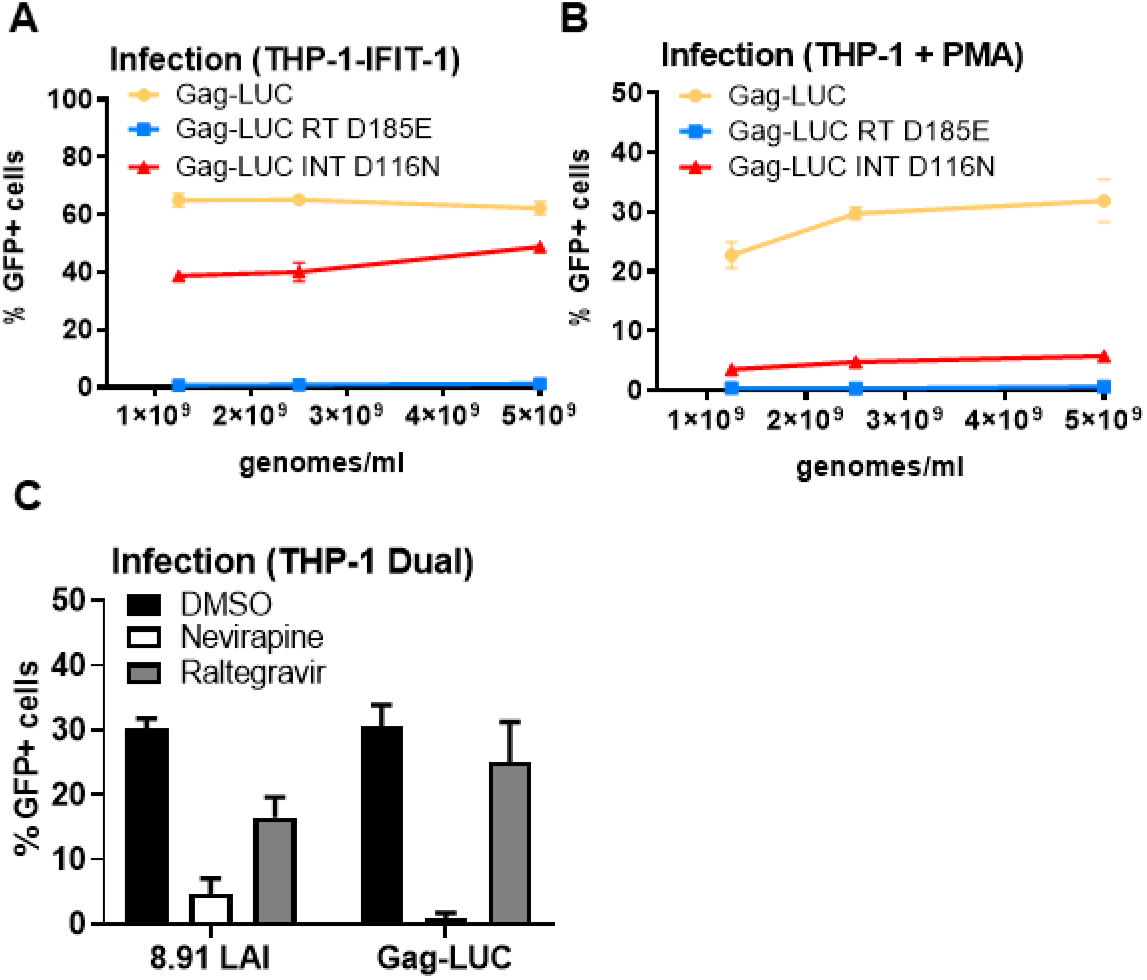
ISG induction by HIV-1 Gag-fusion virus is RT-dependent A: Infection data from Fig 3A. THP-1-IFIT-1 cells transduced for 48 h with Gag-LUC, RT-defective Gag-LUC (Gag-LUC RT D185E) or integrase-defective Gag-LUC (Gag-LUC INT D116N) at 1.25×10^9^, 2.5×10^9^ and 5×10^9^ genomes/ml. B: Infection data from Fig 3B-D. PMA-treated THP-1 shSAMHD1 cells transduced for 48 h with Gag-LUC, Gag-LUC RT D185E or Gag-LUC INT D116N at 1.25×10^9^, 2.5×10^9^ and 5×10^9^ genomes/ml. C: Infection data from Fig 3E-G. THP-1 Dual cells transduced for 48 h with 8.91 LAI or Gag- Luc (1.5 U RT/ml) in the presence of DMSO vehicle, 5 μM neviripine or 10 μM raltegravir. Data are mean ± SD, n = 3, representative of at least 3 repeats.

**Suppl. Fig 4:**
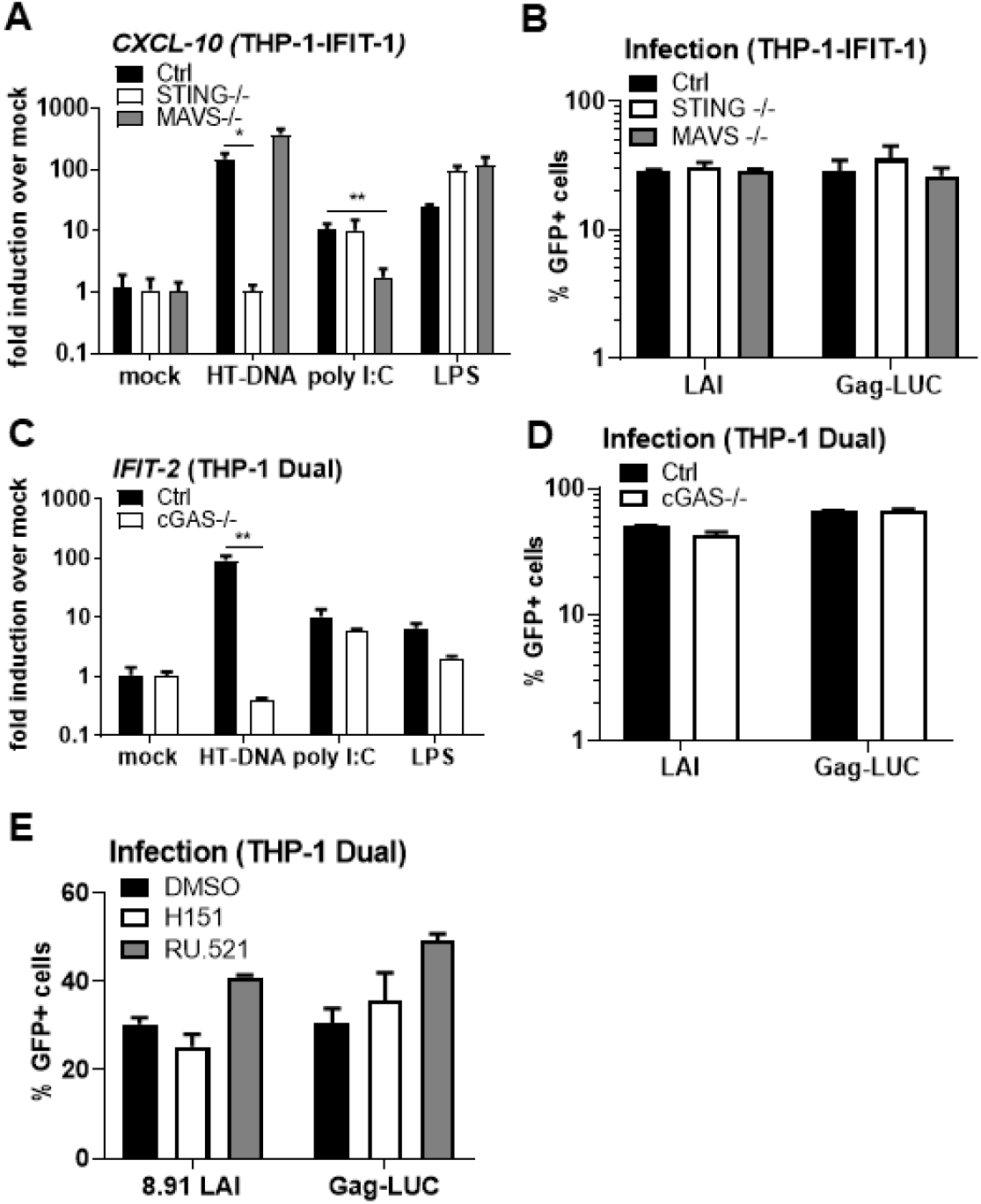
ISG induction by HIV-1 Gag-fusion virus is dependent on cGAS and STING A: *CXCL-10* ISG qPCR from monocytic THP-1-IFIT-1 cells lacking STING or MAVS, or a gRNA control (Ctrl) cell line stimulated for 24 h with 0.1 μg/ml HT-DNA, 0.5 μg/ml poly I:C or 50 ng/ml LPS. B: Infection data from Fig 4A-C. THP-1-IFIT-1 cells lacking STING or MAVS, or a gRNA control (Ctrl) cell line transduced for 48 h with WT LAI or Gag-LUC (1.5 U RT/ml). C: *IFIT-2* ISG qPCR from monocytic THP-1 Dual cells lacking cGAS, or a gRNA control (Ctrl) cell line stimulated for 24 h with 0.1 μg/ml HT-DNA, 0.5 μg/ml poly I:C or 50 ng/ml LPS. D: Infection data from Fig 4D-F. THP-1 Dual cells lacking cGAS, or a gRNA control (Ctrl) cell line transduced for 48 h with WT LAI or Gag-LUC (1.5 U RT/ml). E: Infection data from Fig 4G. THP-1 Dual cells lacking cGAS, or a gRNA control (Ctrl) cell line transduced for 48 h with WT LAI or Gag-LUC (1.5 U RT/ml) in the presence of DMSO vehicle, 0.5 μg/ml STING inhibitor H151 or 10 μg/ml cGAS inhibitor RU.521 Data are mean ± SD, n = 3, representative of at least 3 repeats. Statistical analyses were performed using Student’s t-test, with Welch’s correction where appropriate, comparing to Ctrl cells as indicated. *P < 0.05, **P < 0.01.

**Suppl. Fig 5:**
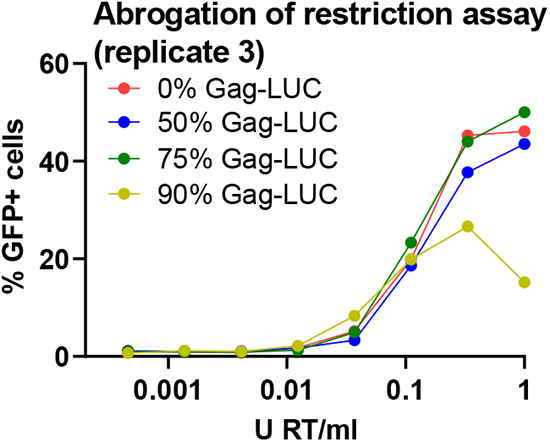
Gag-fusion viruses have reduced capacity to saturate TRIM5α Third replicate assay of data presented in Fig. 5B. FRhK4 cells were co-transduced with a fixed dose of WT LAI.GFP (5×10^7^ genomes/ml) and increasing doses of the WT/Gag-LUC chimeric viruses carrying a luciferase-expressing genome (0.0005 – 1 U RT/ml). Rescue of GFP infectivity was assessed by flow cytometry at 48 h. Data are singlet % GFP values.

## References

1. Chow J, Franz KM, Kagan JC: PRRs are watching you: Localization of innate sensing and signaling regulators. Virology 2015, 479–480:104-109.

2. Schneider WM, Chevillotte MD, Rice CM: Interferon-stimulated genes: a complex web of host defenses. Annu Rev Immunol 2014, 32:513–545.

3. Gringhuis SI, Hertoghs N, Kaptein TM, Zijlstra-Willems EM, Sarrami-Forooshani R, Sprokholt JK, van Teijlingen NH, Kootstra NA, Booiman T, van Dort KA, et al: HIV-1 blocks the signaling adaptor MAVS to evade antiviral host defense after sensing of abortive HIV-1 RNA by the host helicase DDX3. Nat Immunol 2017, 18:225–235.

4. Ringeard M, Marchand V, Decroly E, Motorin Y, Bennasser Y: FTSJ3 is an RNA 2’- O-methyltransferase recruited by HIV to avoid innate immune sensing. Nature 2019, 565:500–504.

5. Gao D, Wu J, Wu YT, Du F, Aroh C, Yan N, Sun L, Chen ZJ: Cyclic GMP-AMP synthase is an innate immune sensor of HIV and other retroviruses. Science 2013, 341:903–906.

6. Lahaye X, Satoh T, Gentili M, Cerboni S, Conrad C, Hurbain I, El Marjou A, Lacabaratz C, Lelievre JD, Manel N: The capsids of HIV-1 and HIV-2 determine immune detection of the viral cDNA by the innate sensor cGAS in dendritic cells. Immunity 2013, 39:1132–1142.

7. Rasaiyaah J, Tan CP, Fletcher AJ, Price AJ, Blondeau C, Hilditch L, Jacques DA, Selwood DL, James LC, Noursadeghi M, Towers GJ: HIV-1 evades innate immune recognition through specific cofactor recruitment. Nature 2013, 503:402–405.

8. Jakobsen MR, Bak RO, Andersen A, Berg RK, Jensen SB, Tengchuan J, Laustsen A, Hansen K, Ostergaard L, Fitzgerald KA, et al: IFI16 senses DNA forms of the lentiviral replication cycle and controls HIV-1 replication. Proc Natl Acad Sci U S A 2013, 110:E4571–4580.

9. Jonsson KL, Laustsen A, Krapp C, Skipper KA, Thavachelvam K, Hotter D, Egedal JH, Kjolby M, Mohammadi P, Prabakaran T, et al: IFI16 is required for DNA sensing in human macrophages by promoting production and function of cGAMP. Nat Commun 2017, 8:14391.

10. Yoh SM, Schneider M, Seifried J, Soonthornvacharin S, Akleh RE, Olivieri KC, De Jesus PD, Ruan C, de Castro E, Ruiz PA, et al: PQBP1 Is a Proximal Sensor of the cGAS-Dependent Innate Response to HIV-1. Cell 2015, 161:1293–1305.

11. Yoh SM, Mamede JI, Lau D, Ahn N, Sanchez-Aparicio MT, Temple J, Tuckwell A, Fuchs NV, Cianci GC, Riva L, et al: Recognition of HIV-1 capsid by PQBP1 licenses an innate immune sensing of nascent HIV-1 DNA. Mol Cell 2022, 82:2871–2884 e2876.

12. Lahaye X, Gentili M, Silvin A, Conrad C, Picard L, Jouve M, Zueva E, Maurin M, Nadalin F, Knott GJ, et al: NONO Detects the Nuclear HIV Capsid to Promote cGAS-Mediated Innate Immune Activation. Cell 2018, 175:488–501 e422.

13. Stavrou S, Aguilera AN, Blouch K, Ross SR: DDX41 Recognizes RNA/DNA Retroviral Reverse Transcripts and Is Critical for In Vivo Control of Murine Leukemia Virus Infection. mBio 2018, 9.

14. Ablasser A, Goldeck M, Cavlar T, Deimling T, Witte G, Rohl I, Hopfner KP, Ludwig J, Hornung V: cGAS produces a 2’-5’-linked cyclic dinucleotide second messenger that activates STING. Nature 2013, 498:380–384.

15. Sun L, Wu J, Du F, Chen X, Chen ZJ: Cyclic GMP-AMP synthase is a cytosolic DNA sensor that activates the type I interferon pathway. Science 2013, 339:786–791.

16. Wu J, Sun L, Chen X, Du F, Shi H, Chen C, Chen ZJ: Cyclic GMP-AMP is an endogenous second messenger in innate immune signaling by cytosolic DNA. Science 2013, 339:826–830.

17. Tanaka Y, Chen ZJ: STING specifies IRF3 phosphorylation by TBK1 in the cytosolic DNA signaling pathway. Sci Signal 2012, 5:ra20.

18. Liu S, Cai X, Wu J, Cong Q, Chen X, Li T, Du F, Ren J, Wu YT, Grishin NV, Chen ZJ: Phosphorylation of innate immune adaptor proteins MAVS, STING, and TRIF induces IRF3 activation. Science 2015, 347:aaa2630.

19. Shang G, Zhang C, Chen ZJ, Bai XC, Zhang X: Cryo-EM structures of STING reveal its mechanism of activation by cyclic GMP-AMP. Nature 2019, 567:389–393.

20. Ishikawa H, Barber GN: STING is an endoplasmic reticulum adaptor that facilitates innate immune signalling. Nature 2008, 455:674–678.

21. Colomer-Lluch M, Ruiz A, Moris A, Prado JG: Restriction Factors: From Intrinsic Viral Restriction to Shaping Cellular Immunity Against HIV-1. Front Immunol 2018, 9:2876.

22. Manel N, Hogstad B, Wang Y, Levy DE, Unutmaz D, Littman DR: A cryptic sensor for HIV-1 activates antiviral innate immunity in dendritic cells. Nature 2010, 467:214–217.

23. Cingoz O, Goff SP: HIV-1 Is a Poor Inducer of Innate Immune Responses. MBio 2019, 10.

24. Tsang J, Chain BM, Miller RF, Webb BL, Barclay W, Towers GJ, Katz DR, Noursadeghi M: HIV-1 infection of macrophages is dependent on evasion of innate immune cellular activation. AIDS 2009, 23:2255–2263.

25. Zuliani-Alvarez L, Govasli ML, Rasaiyaah J, Monit C, Perry SO, Sumner RP, McAlpine-Scott S, Dickson C, Rifat Faysal KM, Hilditch L, et al: Evasion of cGAS and TRIM5 defines pandemic HIV. Nat Microbiol 2022, 7:1762–1776.

26. Sumner RP, Harrison L, Touizer E, Peacock TP, Spencer M, Zuliani-Alvarez L, Towers GJ: Disrupting HIV-1 capsid formation causes cGAS sensing of viral DNA. EMBO J 2020, 39:e103958.

27. Zila V, Margiotta E, Turonova B, Muller TG, Zimmerli CE, Mattei S, Allegretti M, Borner K, Rada J, Muller B, et al: Cone-shaped HIV-1 capsids are transported through intact nuclear pores. Cell 2021, 184:1032–1046 e1018.

28. Muller TG, Zila V, Peters K, Schifferdecker S, Stanic M, Lucic B, Laketa V, Lusic M, Muller B, Krausslich HG: HIV-1 uncoating by release of viral cDNA from capsid-like structures in the nucleus of infected cells. Elife 2021, 10.

29. Li C, Burdick RC, Nagashima K, Hu WS, Pathak VK: HIV-1 cores retain their integrity until minutes before uncoating in the nucleus. Proc Natl Acad Sci U S A 2021, 118.

30. Shen Q, Kumari S, Xu C, Jang S, Shi J, Burdick RC, Levintov L, Xiong Q, Wu C, Devarkar SC, et al: The capsid lattice engages a bipartite NUP153 motif to mediate nuclear entry of HIV-1 cores. Proc Natl Acad Sci U S A 2023, 120:e2202815120.

31. Kim K, Dauphin A, Komurlu S, McCauley SM, Yurkovetskiy L, Carbone C, Diehl WE, Strambio-De-Castillia C, Campbell EM, Luban J: Cyclophilin A protects HIV-1 from restriction by human TRIM5alpha. Nat Microbiol 2019, 4:2044–2051.

32. Peden K, Emerman M, Montagnier L: Changes in growth properties on passage in tissue culture of viruses derived from infectious molecular clones of HIV-1LAI, HIV-1MAL, and HIV-1ELI. Virology 1991, 185:661–672.

33. Mankan AK, Schmidt T, Chauhan D, Goldeck M, Honing K, Gaidt M, Kubarenko AV, Andreeva L, Hopfner KP, Hornung V: Cytosolic RNA:DNA hybrids activate the cGAS-STING axis. EMBO J 2014, 33:2937–2946.

34. Tie CH, Fernandes L, Conde L, Robbez-Masson L, Sumner RP, Peacock T, Rodriguez-Plata MT, Mickute G, Gifford R, Towers GJ, et al: KAP1 regulates endogenous retroviruses in adult human cells and contributes to innate immune control. EMBO Rep 2018, 19.

35. Quintas-Cardama A, Vaddi K, Liu P, Manshouri T, Li J, Scherle PA, Caulder E, Wen X, Li Y, Waeltz P, et al: Preclinical characterization of the selective JAK1/2 inhibitor INCB018424: therapeutic implications for the treatment of myeloproliferative neoplasms. Blood 2010, 115:3109–3117.

36. Bonczkowski P, De Scheerder MA, Malatinkova E, Borch A, Melkova Z, Koenig R, De Spiegelaere W, Vandekerckhove L: Protein expression from unintegrated HIV-1 DNA introduces bias in primary in vitro post-integration latency models. Sci Rep 2016, 6:38329.

37. Van Loock M, Hombrouck A, Jacobs T, Winters B, Meersseman G, Van Acker K, Clayton RF, Malcolm BA: Reporter gene expression from LTR-circles as tool to identify HIV-1 integrase inhibitors. J Virol Methods 2013, 187:238–247.

38. Haag SM, Gulen MF, Reymond L, Gibelin A, Abrami L, Decout A, Heymann M, van der Goot FG, Turcatti G, Behrendt R, Ablasser A: Targeting STING with covalent small-molecule inhibitors. Nature 2018, 559:269–273.

39. Vincent J, Adura C, Gao P, Luz A, Lama L, Asano Y, Okamoto R, Imaeda T, Aida J, Rothamel K, et al: Small molecule inhibition of cGAS reduces interferon expression in primary macrophages from autoimmune mice. Nat Commun 2017, 8:750.

40. Ganser-Pornillos BK, Chandrasekaran V, Pornillos O, Sodroski JG, Sundquist WI, Yeager M: Hexagonal assembly of a restricting TRIM5alpha protein. Proc Natl Acad Sci U S A 2011, 108:534–539.

41. Li YL, Chandrasekaran V, Carter SD, Woodward CL, Christensen DE, Dryden KA, Pornillos O, Yeager M, Ganser-Pornillos BK, Jensen GJ, Sundquist WI: Primate TRIM5 proteins form hexagonal nets on HIV-1 capsids. Elife 2016, 5.

42. Fletcher AJ, Christensen DE, Nelson C, Tan CP, Schaller T, Lehner PJ, Sundquist WI, Towers GJ: TRIM5alpha requires Ube2W to anchor Lys63-linked ubiquitin chains and restrict reverse transcription. EMBO J 2015, 34:2078–2095.

43. Fletcher AJ, Vaysburd M, Maslen S, Zeng J, Skehel JM, Towers GJ, James LC: Trivalent RING Assembly on Retroviral Capsids Activates TRIM5 Ubiquitination and Innate Immune Signaling. Cell Host Microbe 2018, 24:761–775 e766.

44. Pertel T, Hausmann S, Morger D, Zuger S, Guerra J, Lascano J, Reinhard C, Santoni FA, Uchil PD, Chatel L, et al: TRIM5 is an innate immune sensor for the retrovirus capsid lattice. Nature 2011, 472:361–365.

45. Jacques DA, McEwan WA, Hilditch L, Price AJ, Towers GJ, James LC: HIV-1 uses dynamic capsid pores to import nucleotides and fuel encapsidated DNA synthesis. Nature 2016, 536:349–353.

46. Shi J, Aiken C: Saturation of TRIM5 alpha-mediated restriction of HIV-1 infection depends on the stability of the incoming viral capsid. Virology 2006, 350:493–500.

47. Khan H, Sumner RP, Rasaiyaah J, Tan CP, Rodriguez-Plata MT, Van Tulleken C, Fink D, Zuliani-Alvarez L, Thorne L, Stirling D, et al: HIV-1 Vpr antagonizes innate immune activation by targeting karyopherin-mediated NF-kappaB/IRF3 nuclear transport. Elife 2020, 9.

48. Okumura A, Alce T, Lubyova B, Ezelle H, Strebel K, Pitha PM: HIV-1 accessory proteins VPR and Vif modulate antiviral response by targeting IRF-3 for degradation. Virology 2008, 373:85–97.

49. Trotard M, Tsopoulidis N, Tibroni N, Willemsen J, Binder M, Ruggieri A, Fackler OT: Sensing of HIV-1 Infection in Tzm-bl Cells with Reconstituted Expression of STING. J Virol 2016, 90:2064–2076.

50. Vermeire J, Roesch F, Sauter D, Rua R, Hotter D, Van Nuffel A, Vanderstraeten H, Naessens E, Iannucci V, Landi A, et al: HIV Triggers a cGAS-Dependent, Vpu- and Vpr-Regulated Type I Interferon Response in CD4(+) T Cells. Cell Rep 2016, 17:413–424.

51. Sauter D, Hotter D, Van Driessche B, Sturzel CM, Kluge SF, Wildum S, Yu H, Baumann B, Wirth T, Plantier JC, et al: Differential regulation of NF-kappaB-mediated proviral and antiviral host gene expression by primate lentiviral Nef and Vpu proteins. Cell Rep 2015, 10:586–599.

52. Sumner RP, Thorne LG, Fink DL, Khan H, Milne RS, Towers GJ: Are Evolution and the Intracellular Innate Immune System Key Determinants in HIV Transmission? Front Immunol 2017, 8:1246.

53. Yan N, Regalado-Magdos AD, Stiggelbout B, Lee-Kirsch MA, Lieberman J: The cytosolic exonuclease TREX1 inhibits the innate immune response to human immunodeficiency virus type 1. Nat Immunol 2010, 11:1005–1013.

54. Siddiqui MA, Saito A, Halambage UD, Ferhadian D, Fischer DK, Francis AC, Melikyan GB, Ambrose Z, Aiken C, Yamashita M: A Novel Phenotype Links HIV-1 Capsid Stability to cGAS-Mediated DNA Sensing. J Virol 2019, 93.

55. Renner N, Kleinpeter A, Mallery DL, Albecka A, Rifat Faysal KM, Bocking T, Saiardi A, Freed EO, James LC: HIV-1 is dependent on its immature lattice to recruit IP6 for mature capsid assembly. Nat Struct Mol Biol 2023.

56. Civril F, Deimling T, de Oliveira Mann CC, Ablasser A, Moldt M, Witte G, Hornung V, Hopfner KP: Structural mechanism of cytosolic DNA sensing by cGAS. Nature 2013, 498:332–337.

57. Yant SR, Mulato A, Hansen D, Tse WC, Niedziela-Majka A, Zhang JR, Stepan GJ, Jin D, Wong MH, Perreira JM, et al: A highly potent long-acting small-molecule HIV-1 capsid inhibitor with efficacy in a humanized mouse model. Nat Med 2019, 25:1377–1384.

58. Marquez CL, Lau D, Walsh J, Shah V, McGuinness C, Wong A, Aggarwal A, Parker MW, Jacques DA, Turville S, Bocking T: Kinetics of HIV-1 capsid uncoating revealed by single-molecule analysis. Elife 2018, 7.

59. Kanamoto T, Kashiwada Y, Kanbara K, Gotoh K, Yoshimori M, Goto T, Sano K, Nakashima H: Anti-human immunodeficiency virus activity of YK-FH312 (a betulinic acid derivative), a novel compound blocking viral maturation. Antimicrob Agents Chemother 2001, 45:1225–1230.

60. Zufferey R, Nagy D, Mandel RJ, Naldini L, Trono D: Multiply attenuated lentiviral vector achieves efficient gene delivery in vivo. Nat Biotechnol 1997, 15:871–875.

61. Vermeire J, Naessens E, Vanderstraeten H, Landi A, Iannucci V, Van Nuffel A, Taghon T, Pizzato M, Verhasselt B: Quantification of reverse transcriptase activity by real-time PCR as a fast and accurate method for titration of HIV, lenti- and retroviral vectors. PLoS One 2012, 7:e50859.

